# FDA-approved PDE4 inhibitors reduce the dominant toxicity of ALS–FTD-associated *CHCHD10^S59L^*

**DOI:** 10.1101/2024.06.04.597429

**Authors:** Swati Maitra, Minwoo Baek, Yun-Jeong Choe, Nam Chul Kim

## Abstract

Mutations in coiled-coil-helix-coiled-coil-helix domain containing 10(*CHCHD10*) have been identified as a genetic cause of amyotrophic lateral sclerosis and/or frontotemporal dementia(ALS-FTD). In our previous studies using in vivo *Drosophila* model expressing C2C10H^S81L^, and human cell models expressing CHCHD10^S59L^, we have identified that the PINK1/Parkin pathway is activated and causes cellular toxicity. Furthermore, we demonstrated that pseudo-substrate inhibitors for PINK1 and mitofusin2 agonists mitigated the cellular toxicity of CHCHD10^S59L^. Evidences using *in vitro/ in vivo* genetic and chemical tools indicate that inhibiting PINK1 would be the most promising treatment for CHCHD10^S59L^-induced diseases. Therefore, we have investigated cellular pathways that can modulate the PINK1/Parkin pathway and reduce CHCHD10^S59L^-induced cytotoxicity. Here, we report that FDA-approved PDE4 inhibitors reduced CHCHD10^S59L^-induced morphological and functional mitochondrial defects in human cells and an *in vivo Drosophila* model expressing C2C10H^S81L^. Multiple PDE4 inhibitors decreased PINK1 accumulation and downstream mitophagy induced by CHCHD10^S59L^. These findings suggest that PDE4 inhibitors currently available in the market may be repositioned to treat CHCHD10^S59L^-induced ALS-FTD and possibly other related diseases.

## INTRODUCTION

Coiled-coil-helix-coiled-coil-helix domain containing 10(*CHCHD10*) encodes a mitochondrial protein that causes amyotrophic lateral sclerosis and/or frontotemporal dementia(ALS-FTD) when it is mutated(Bannwarth et al., 2014). We reported that CHCHD10^S59L^ overexpression in human cells induced fragmented mitochondria and mitochondrial dysfunction. Consistently, expressing C2C10H^S81L^(*Drosophila* homolog of human CHCHD2 and CHCHD10 bearing a mutation in the conserved postion with human S59L) with the tissue-specific GAL4/UAS expression system, induced rough eye phenotypes, neuromuscular junction defects, and muscle degeneration with fragmented mitochondria(Baek et al., 2021). Through a series of genetic interaction studies, we identified that PINK1 and Parkin are strong modifiers for the dominant toxicity of C2C10H^S81L^. RNAi-mediated knockdown of PINK1 and Parkin strongly suppressed the rough eye phenotypes, muscle degeneration, and mitochondrial fragmentation of C2C10H^S81L^. Additionally, ATP production was recovered in the adult indirect flight muscles. In contrast, overexpression of PINK1 and Parkin significantly enhanced the rough eye phenotypes(Baek et al., 2021).

CHCHD10^S59L^ expression in human cells including HeLa and SH-SY5Y induced mitochondrial accumulation of PINK1 and Parkin, and subsequently, LC3 which is a marker for activated mitophagy(Baek et al., 2021). siRNA transfection for PINK1 and Parkin reduced mitochondrial fragmentation induced by CHCHD10^S59L^ and also improved functional mitochondrial respiration significantly. When we knocked-down mitophagy adaptors for the PINK1/Parkin pathway, it showed recovery in functional mitochondrial respiration in multiple cell lines. Interestingly, CHCHD10^S59L^ induced mitochondrial fragmentation and dysfunction in HeLa that does not express Parkin. siRNA mediated knock-down of PINK1 reversed the mitochondrial network fragmentation in two patient-derived fibroblasts(Baek et al., 2021). Therefore, these results indicate that reducing PINK1 expression or activity can be a potential therapeutic strategy.

In fact, we identified two psudosubstrate inhibitors for PINK1 and demonstrated that two peptide inhibitors reduced its ubiquitin kinase activity and mitigated CHCHD10^S59L^-induced mitochondria fragmentation(Baek et al., 2021). However, due to the lack of small molecule PINK1 inhibitors, we could not evaluate the efficacy of PINK1 inhibition *in vivo*. Therefore, we searched small molelcules that may reduce PINK1 expression level or enzymatic activity directly or indirectly.

In 2016, Akabane et al. reported that PKA regulates the stability of PINK1 via phosphorylating mitofilin(Akabane et al., 2016). In that study, they demonstrated that Parkin recruitment to depolarized mitochondria was severely inhibited by Forskolin treatment via increase in cAMP levels and activation of a protein kinase A(PKA). Activation of PKA and subsequent phosphorylation of MIC60(which is a substrate of PKA) impaired the translocation of Parkin to damaged mitochondria via reduced PINK1 protein stability. Therefore, we evaluated the efficacy of PDE4 inhibitors for ameliorating CHCHD10^S59L^-induced toxicity in human cells and *Drosophila*. Here, we report that FDA-approved PDE4 inhibitors mitigate CHCHD10^S59L^(C2C10H^S81L^)-induced toxicity in human cells and *Drosophila*.

## RESULTS

### Forskolin blocks PINK1 and Parkin accumulation

To validate the results from Akabane et al.’s research, we tested whether an adenylate cyclase activator, Forskolin, reduces PINK1 and Parkin accumulation caused by CCCP. CCCP is a mitochondrial uncoupling agent inducing PINK1 stabilization and subsequent Parkin accumulation on mitochondria. As they demonstrated, acute Forskolin(100uM) treatment almost completely blocked Parkin accumulation induced by CCCP in HeLa cells stably expressing Parkin-YFP(Fig. 1A-B). As expected, we also successfully detected that Forskolin treatment blocked PINK1 accumulation significantly(Fig. 1C-D). These observations led us to examine the effects of Forskolin on CHCHD10^S59L^-induced mitochondrial toxicity.

**Figure 1.**
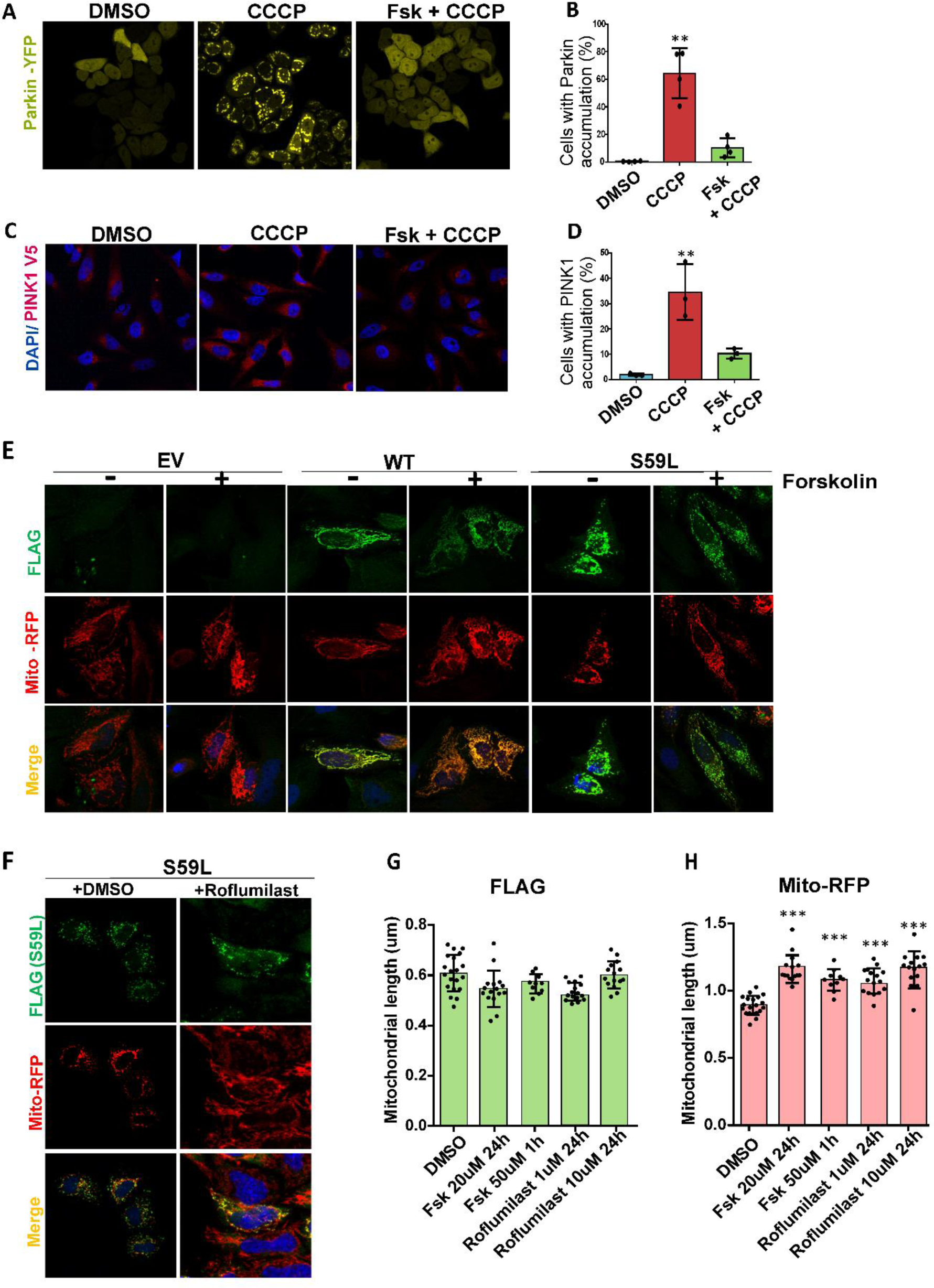
Forskolin blocks PINK1-Parkin accumulation and mitigates CHCHD10^S59L^ induced cellular toxicity. HeLa ^YFP-Parkin^cells and HeLa ^PINK1-V5-His^cells were pre-treated with Forskolin(100uM-1hr) followed by CCCP(10uM-2hrs). **(A)**Immunohistochemical images for visualization mitochondrial Parkin-YFP accumulation in DMSO, CCCP, Fsk+CCCP groups respectively. **(B)**Graphical representation indicating percentage of mitochondrial Parkin accumulation in cells. **(C)**Immunohistochemical images for visualization of PINK1 accumulation in cells immunostained with anti-V5 antibody(red). **(D)**A representative graph indicating percentage PINK1 accumulation. Data are shown as mean ± SD (one-way ANOVA with post hoc Dunnett’s Test); n = 4(Parkin), n=3(PINK1) independent experiments >200 cells for each group). **(E)**HeLa cells were transfected with Empty Vector, FLAG-tagged CHCHD10^WT^, CHCHD10^S59L^ in the DMSO, Forskolin containing medium. Cells were infected with Mito-RFP (Red, mitochondria) for overnight. Representative images of cells immunostained with anti-FLAG antibody (green, CHCHD10^S59L^), Mito-RFP and DAPI. **(F)**Representative images of CHCHD10^S59L^ expressing mitochondria cultured in DMSO, and Roflumilast containing media. **(G)(H)**Quantification of CHCHD10-FLAG (green) and Mito-RFP (red) signal length. Data are shown as mean ± SD (one-way ANOVA and post hoc Dunnett test, two-sided comparison with DMSO).

### Forskolin and Roflumilast mitigate the toxicity of CHCHD10^S59L^ in cell models

When we treated Forskolin to HeLa cells transiently transfected with CHCHD10^S59L^, mitochondrial branch length increased significantly compared to that of DMSO-treated cells(Fig. 1E). Since Forskolin increases cyclic AMP levels by activating adenylate cyclases, we investigated whether phosphodiesterase inhibitors(PDE inhibitors) could induce the same effect by preventing cAMP degradation. First, we chose Roflumilast, a selective PDE4 inhibitor, that was approved and marketed for chronic obstructive pulmonary disease(COPD)(Wedzicha et al., 2016) (Kelly et al., 2017). When we treated Roflumilast(1 µM or 10 µM for 24hrs) to CHCHD10^S59L^-expressing HeLa cells, mitochondrial fragmentation caused by CHCHD10^S59L^ was significantly reduced while aggregation of CHCHD10^S59L^ was not affected(Fig. 1F,G,H).

### Forskolin and Roflumilast reduce PINK1 accumulation caused by CHCHD10^S59L^ in HeLa^PINK1-V5^ cells

We previously reported that CHCHD10^S59L^ induced PINK1 accumulation on mitochondria and reduction of PINK1 expression or activity was beneficial to mitochondria morphologically and functionally(Baek et al., 2021). Therefore, we examined whether Forskolin and Roflumilast reduced CHCHD10^S59L^-induced PINK1 acculumation in HeLa cells. For this purpose, we used HeLa cells stably expressing V5 tagged PINK1(HeLa^PINK1-V5^ cells)(Narendra et al., 2010). When CHCHD10^S59L^ was transfected and expressed, more than 80% of cells showed PINK1 accumulation on mitochondria and the PINK1 aggregation-positive cells were significantly decreased by Forskolin and Roflumilast treatment(Fig. 2A-B). Acute treatment of 50 µM Forskolin and 24 hour-treatment of 20 µM Forskolin showed similar effects(Fig. 2B). However, we did not observe significant dose-dependent effects of Roflumilast when it was treated with 1 µM or 10 µM concentrations in this experiment(Fig 2B).

**Figure 2.**
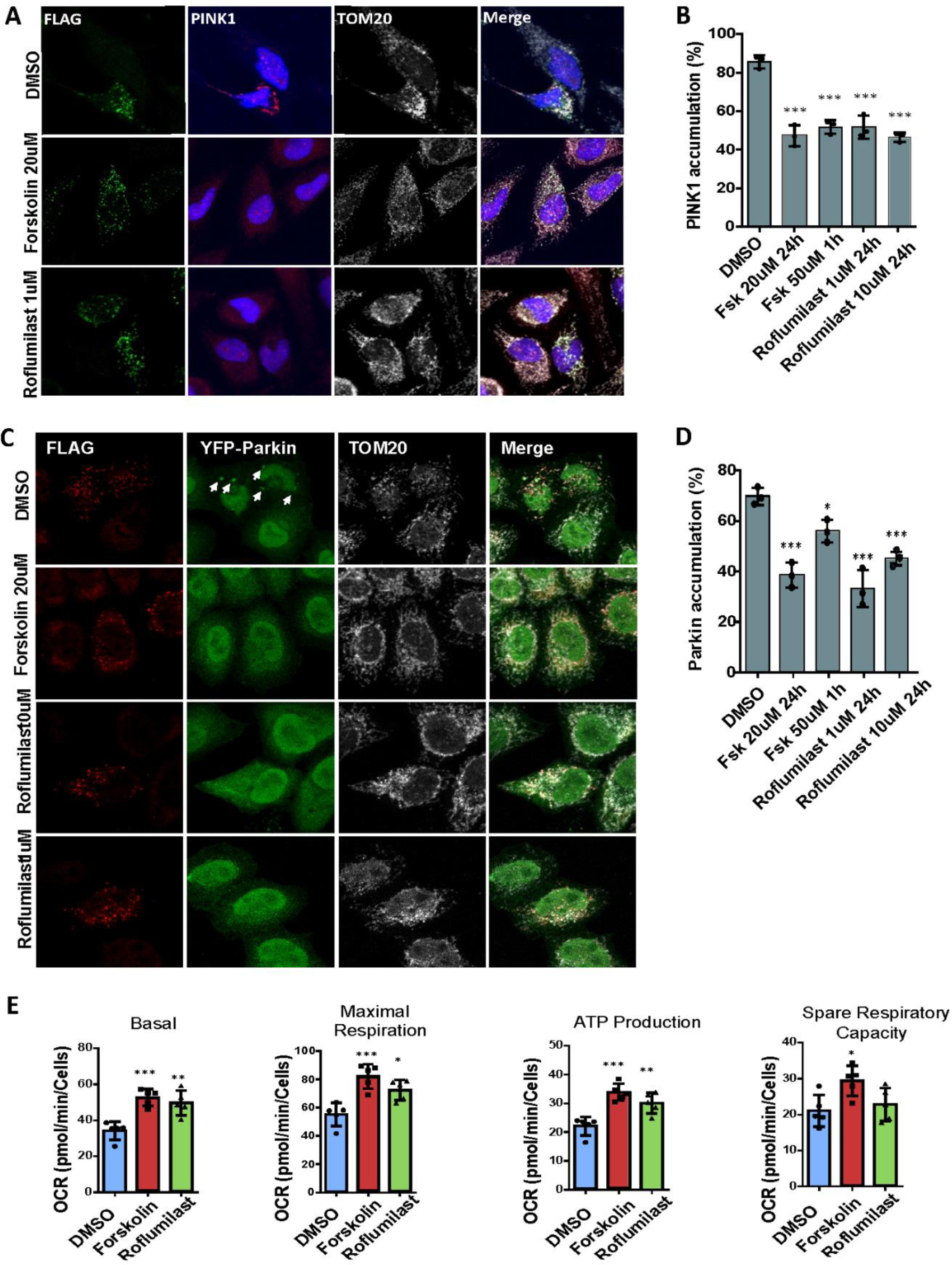
Forskolin and Roflumilast reduce PINK1-Parkin accumulation induced by CHCHD10^S59L^ expression in HeLa cells. HeLa^PINK1-V5^ cells and HeLa^YFP-Parkin^ cells were transfected with FLAG-tagged CHCHD10^S59L^ in the DMSO, Forskolin, and Roflumilast containing medium. **(A)**Representative images of CHCHD10^S59L^ expressing mitochondria cultured in DMSO, Forskolin and Roflumilast containing media. Cells were immunostained with anti-FLAG(green, CHCHD10^S59L^) and anti-PINK1(red, PINK1) and anti-TOM20(white, mitochondria) antibodies. **(B)**Bar graphs indicating percentage of HeLa^PINK1-V5^ showing PINK1 accumulation in CHCHD10^S59L^ expressing mitochondria. Data are shown as mean ± SD (one-way ANOVA and post hoc Dunnett test, two-sided, comparison with DMSO). **(C)**Representative images from CHCHD10^S59L^ expressing mitochondria cultured in DMSO, Forskolin(20 μM) and Roflumilast(1 μM) containing media. Cells were immunostained against FLAG (green, CHCHD10^S59L^) and TOM20 (white, mitochondria) antibodies. Arrows indicate accumulated parkin (green, YFP-parkin) in the mitochondria. **(D)**Bar graphs indicating percentage of HeLa^YFP-Parkin^ showing Parkin accumulation in CHCHD10^S59L^ expressing mitochondria. Data are shown as mean ± SD (one-way ANOVA and post hoc Dunnett test, two-sided, comparison with DMSO. **(E)** Mitochondrial respiration was measured by Seahorse XF Cell Mito Stress tests 24hrs after CHCHD10^S59L^ transfection in DMSO, Forskolin and Roflumilast treated HeLa cells. Data are shown as mean ±SD (one-way ANOVA and post hoc Dunnett test, two-sided, comparison with DMSO, \*\*\**p* < 0.001, \*\**p* < 0.01 and \**p* < 0.05).

### Forskolin and Roflumilast reduce Parkin accumulation caused by CHCHD10^S59L^ in HeLa^Parkin-YFP^ cells

Mitochondria-localized PINK1 phosphorylates ubiquitins and the ubiquitin-like domain of Parkin, resulting in Parkin accumulation on mitochondria. Therefore, we investigated whether Forskolin and Roflumilast reduced Parkin acculumation in HeLa cells via reducing CHCHD10^S59L^-induced PINK1 accumulation. We used HeLa cells stably expressing yellow fluorescent protein(YFP)-tagged Parkin. When CHCHD10^S59L^ was transfected and expressed, about 70% of cells had Parkin-YFP punctae colocalized with mitochondria(Fig. 2C). Although acute 50 µM Forskolin treatment reduced Parkin-YFP accumulation, 24 hour-treatment of 20 µM Forskolin treatment showed a more significant decrease of Parkin accumulation on mitochondria(Fig. 2D). Interestingly, a relatively low dose of Roflumilast(1 µM) significantly reduced Parkin-YFP accumulation more than 10 µM Roflumilast(Fig. 2D). This may be due to the dynamic regulation of cytosolic cAMP levels by cAMP-activated PKA and PKA-activated phosphodiesterase(Bauman et al., 2006). Consistently, the seahorse analysis revealed that Forskolin and/or Roflumilast treatment increased basal mitochondrial respiration and ATP production with increased maximal respiration(Fig. 2E). The spare capacity was also increased minimally and not significantly in Roflumilast treated samples(Fig. 2E). The reduction of Parkin accumulation was also observed in a neuroblastoma cell line, SH-SY5Y, with transiently transfected mCherry-Parkin(Suppl Fig S1 A-B).

### Forskolin and Roflumilast reduced LC3 accumulation on mitochondria

The PINK1/Parkin pathway regulates mitophagy(Jin & Youle, 2012). We have demonstrated that CHCHD10^S59L^ expression led to increased LC3 accumulation and subsequent mitophagy(Baek et al., 2021). Thus, we investigated whether Forskolin and Roflumilast could reduce LC3 accumulation caused by CHCHD10^S59L^. We observed that almost 80% of the CHCHD10^S59L^ transfected HeLa^YFP-Parkin^cells showed LC3 accumulation and Forskolin and Roflumilast treatment reduced the LC3 accumulation significantly(Fig 3A-B).

**Figure 3.**
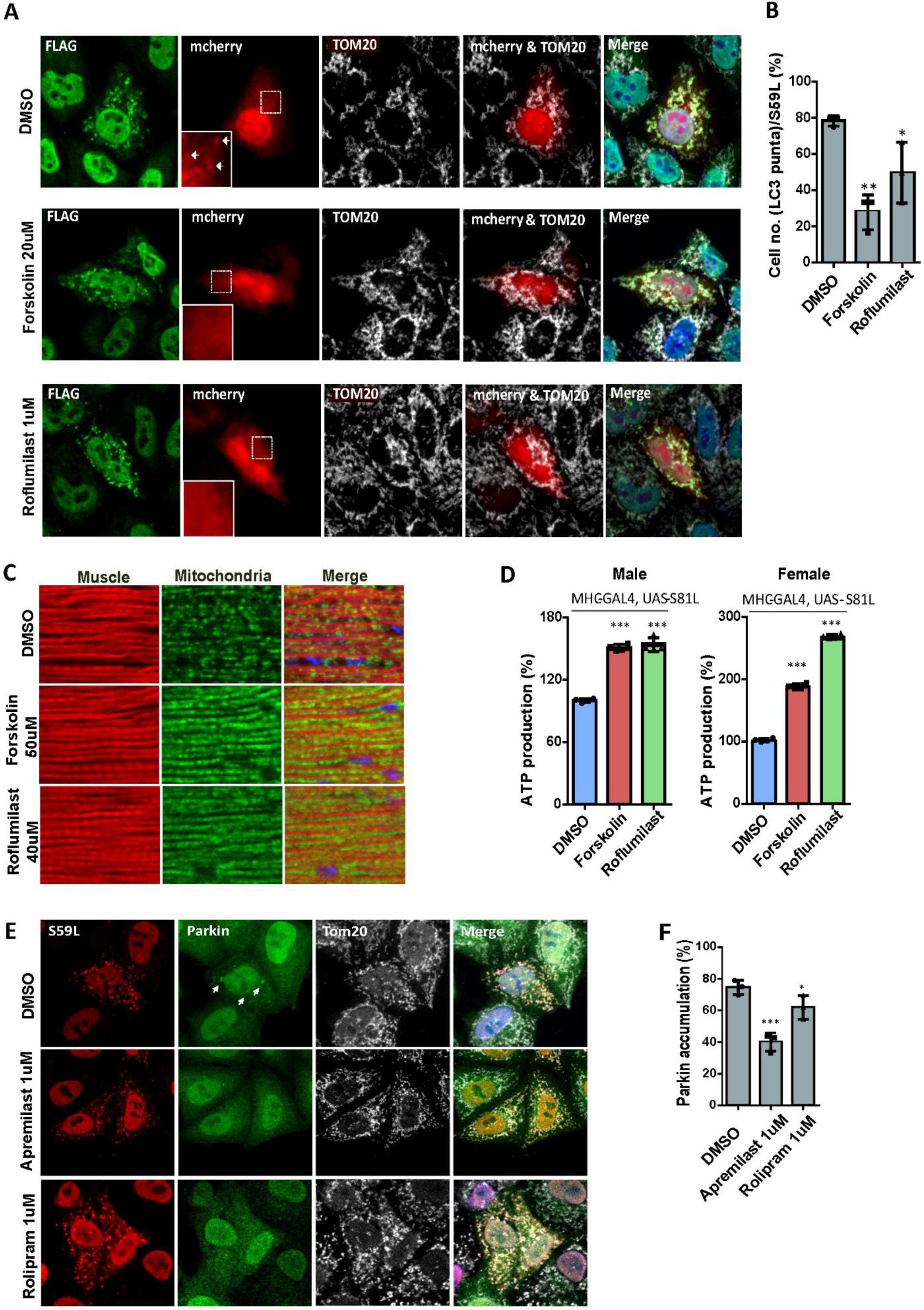
Forskolin and Roflumilast alleviates mitophagy and muscle damage in *Drosophila* and effects of other PDE4 inhibitors. HeLa^YFP-Parkin^cells were co-transfected with FLAG-tagged CHCHD10^S59L^ and mCherry-LC3 in the DMSO, Forskolin, and Roflumilast containing medium. **(A)**Representative images of CHCHD10^S59L^ expressing mitochondria cultured in DMSO, Forskolin(20 μM) and Roflumilast(1 μM) containing media. Cells were immunostained with anti-FLAG (green, CHCHD10^S59L^) and anti-TOM20 (white, mitochondria) antibodies. Arrows indicates LC3 puncta in the CHCHD10^S59L^ expressing mitochondria. **(B)**Percentage of HeLa^YFP-parkin^ showing LC3-mcherry puncta form in the CHCHD10^S59L^ expressing mitochondria. Data are shown as mean ± SD (one-way ANOVA and post hoc Dunnett’s test, two-sided, comparison with DMSO). **(C)**Representative micrographs of indirect flight muscles from 10-day-old adult flies (MHC-Gal4, UAS-S81L) fed with Forskolin(50 μM) and Roflumilast(40 μM) immunostained with streptavidin-Alexa Flour 488 and phalloidin-Alexa Fluor 594. DMSO was used as a vehicle. **(D)**Graphical representation of ATP levels in thoraxes from 10-day-old flies. Data are shown as mean ±SD (one-way ANOVA and *post hoc* Dunnett test, two-sided, \*\*\**p* < 0.001, compared to DMSO. HeLa cells were transfected with FLAG-tagged CHCHD10^S59L^ in the DMSO, Apremilast, and Rolipram containing medium. **(E)**Representative images from CHCHD10^S59L^ expressing mitochondria cultured in DMSO, Apremilast(1 μM) and Rolipram(1 μM) contained media in HeLa cells. Cells were immunostained against anti-FLAG (red, CHCHD10^S59L^) and anti-TOM20 (white, mitochondria) antibodies. Arrows indicates accumulated parkin in the mitochondria. **(F)**Percentage of HeLa showing parkin-YFP accumulation in CHCHD10^S59L^ expressing mitochondria. Data are shown as mean ±SD (one-way ANOVA and post hoc Dunnett test, two-sided, comparison with DMSO, \*\*\**p* < 0.001, \*\*\**p* < 0.01 and \**p* < 0.05).

### Forskolin and Roflumilast ameliorated mitochondrial defects induced by C2C10H^S81L^ in *Drosophila*

Finally, we investigated the effect of Forskolin and Roflumilast treatment on mitochondrial defects in *Drosophila* expressing C2C10H^S81L^. For this, we expressed C2C10H^S81L^ in muscle tissues with the MHC-GAL4 driver. The C2C10H^S81L^ flies were reared in fly food containing Forskolin(50 uM), Roflumilast(40 uM), and DMSO. Indirect flight muscle sections of C2C10H^S81L^ flies stained with a fluorophore conjugated streptavidin(green) and phalloidin(red) revealed fragmented mitochondria along with muscular degeneration(Fig. 3C, DMSO). This condition was ameliorated significantly in Forskolin and Roflumilast treated flies(Fig. 3C). ATP levels were measured as an indicator of mitochondrial dysfunction in the fly thoraxes which primarily consists of muscle tissues. We observed significantly improved ATP levels in the muscle tissues of flies expressing C2C10H^S81L^ treated with Forskolin and Roflumilast, as compared to DMSO-fed C2C10H^S81L^ flies(Fig. 3D). Therefore, Forskolin and Roflumilast imparted a rescue effect on the mitochondrial defects induced by toxic C2C10H^S81L^.

### Other PDE4 inhibitors, Apremilast and Rolipram, are also effective

To further validate the potential of PDE4 inhibitors against CHCHD10^S59L^mediated toxicity, we also tested the first generation PDE4 inhibitor, Rolipram(Kim et al., 2017; Sommer\ et al., 1995), and another approved drug, Apremilast, which is prescribed for plaque psoriasis or psoriatic arthritis(Allawh et al., 2020; McCann et al., 2010). Both Apremilast and Rolipram reduced Parkin-YFP accumulation in HeLa^YFP-Parkin^ cells transiently transfected with CHCHD10^S59L^(Fig. 3E). 1 µM Apremilast reduced Parkin-YFP accumulation to a comparable level with 20µM Roflumilast. However, Rolipram only showed marginal efficacy though it was statistically significant in multiple independently repeated experiments(Fig. 3F). The seahorse analysis also verified that both Apremilast and Rolipram reduced the toxicity of CHCHD10^S59L^ while Apremilast was more effective than Rolipram in rescuing defective mitochondrial function caused by CHCHD10^S59L^(Fig S2 A-C). A similar trend in mitochondrial respiration was also reflected in the seahorse assay using SHSY5Y neuroblastoma cells(Fig S1 C).

## DISCUSSION

Mitochondrial dysfunction remains one of the prominent pathological features of neurodegenartive disorders including ALS-FTD(Zhao et al., 2022). Emerging reports suggested that cAMP signaling plays a key role in mitochondrial biology and homeostasis(Aslam & Ladilov, 2021). Baek and colleagues demonstrated that CHCHD10^S59L^ mutant induced dominant toxicity in *Drosophila* and HeLa cells, and that the PINK1/Parkin mediated pathway was implicated in CHCHD10 associated ALS-FTD(Baek et al., 2021). Here, we demonstrate that FDA-approved PDE4 inhibitors successfully reduced morphological and functional mitochondrial defects in human cell lines and *in vivo Drosophila* model expressing CHCHD10^S59L^ and C2C10H^S81L^, respectively.

We observed that Forskolin treatment reduced the Parkin and PINK1 accumulation in HeLa ^YFP-PARKIN^ and HeLa^PINK1-V5^ cells, respectively, after depolarization by an uncoupling gent as previously reported by Akabane and colleagues(Akabane et al., 2016). Forskolin, being an adenylate cyclase activator, increased the cAMP levels of the system and activated the PKA pathway and led to destabilization of PINK1 on depolarized mitochondria. Forskolin also reduced the disrupted mitochondrial network and enhanced mitochondrial respiration in transiently transfected CHCHD10^S59L^ HeLa cells. Similarly, Roflumilast treatment to CHCHD10^S59L^ expressing cells exhibited a rescued phenotype. Both drugs act through the same mechanism i.e via cAMP/PKA pathway by increasing the cAMP levels in the cellular pool, one by hydrolyzing ATP to cAMP and another by inhibiting the hydrolysis of phosphodiester bond of cAMP to AMP. Following mitochondrial stress, the accumulated PINK1 and Parkin induces mitophagy for the removal of damaged/diseased mitochondria(Pickles et al., 2018). LC3 is an ubiquitin-like protein that is covalently attached to the surface of autophagosome during its biogenesis(Gong et al., 2023). CHCHD10^S59L^ overexpression caused LC3 accumulation in HeLa^YFP-Parkin^ cells as a result of active mitophagy via the PINK1/Parkin pathway. Both Forskolin and Roflumilast treatment led to a decrease in LC3 accumulation in the cells. In our Drosophila model, we also show that Forskolin and Roflumilast reduced the C2C10H^S81L^ -induced toxicity.

PDE4 targeted therapies have shown promising results in various neurological disorders such Alzheimer’s disease (AD), Parkinson’s disease (PD), Fragile X Syndrome, depression and neuropathic pain(Crocetti et al., 2022). At present, very few drugs are available for the treatment of ALS. Ibudilast, a PDE4 inhibitor is in different phases of clinical trials for multiple neurodgenerative diseases, including Multiple Sclerosis and ALS due to their anti-inflammatory activity.In case of ALS, Ibudilast has been shown to slow the disease progression(Oskarsson et al., 2021) (https://alsnewstoday.com/news/als-therapy-ibudilast-safe-well-tolerated-given-intravenously/). Apremilast is also in many clinical trials and crosses the blood brain barrier(Crocetti et al., 2022). Significantly, Apremilast was very effective in mitigating the mitochondrial fragmentation and respiration in lower doses compared to Roflumilast and Forskolin in our experiments. Our data suggest that PDE4 inhibitors may be effective therapeutics in the context of CHCHD10^S59L^-induced ALS-FTD. It also implies that the effectiveness of PDE4 inhibitors for neurodegeneration may not be the only consequence of modulating neuroinflammation as in the case of many previous clinical studies, but also regulating mitochondrial biology(which is the primary mechanism involved in CHCHD10^S59L^ associated ALS-FTD). It is noteworthy that TDP-43 loss increases PDE4 expression(Briese et al., 2020). It may be possible to increase the efficacy of PDE4 inhbitors against neurodegeneration by improving their ability to modulate mitochondrial biology rather than solely focusing on neuroinflammation. It will be interesting to study a combinatorial effect of Forskolin and PDE4 inhibitors on the pathophysiology of ALS-FTD which might act effectively in much lower doses pharmacologically and can be well tolerated in the system without many adverse side effects.

## METHODS

### Chemicals and Drugs

CCCP was purchased from Sigma-Aldrich (C2759). Forskolin was purchased from Millipore-Sigma. PDE4 inhibitors Roflumilast (SML1099-5MG ), Rolipram (R6520-10MG) were bought from Sigma and Apremilast (501872588) was purchased from MedChem Express.

### DNA constructs

All cDNAs for human wild-type and mutant forms of CHCHD10 were synthesized and inserted in the pcDNA3 vector containing FLAG-tag by Genescripts and mCherryLC3B (#40827) plasmid was obtained from Addgene.

### Cell culture and transfections

HeLa cells, HeLa^Parkin-YFP^, HeLa^PINK1-V5^ cells were gifts from Dr. Richard Youle’s lab. Cells were maintained in Dulbecco-modified Eagle medium (Gibco) supplemented with 10% fetal bovine serum (Gibco), 1× penicillin/streptomycin (Invitrogen), and 1 x GlutaMax (Gibco). Cells were transfected using jetPRIME (Polyplus #89129-924) transfection reagent according to the manufacturer’s protocol. For plasmid transfections, around 3 X10^4^ cells were counted using ADAM-MC2 cell counter (NanoEntek) and plated in a 4-well chambered slides in a drug containing media and allowed to grow for 24 hrs. Transfections were carried out with 0.5 ug of plasmid and 1ul of jetPRIME transfection reagent (1:2) mixed in 50uL of jetPRIME buffer per well. In cases of co-transfections, plasmids were mixed at equimolar concentrations. Media was changed with a fresh drug containing media after 4 hrs and incubated with Mito-RFP (CellLight™ Mitochondria-RFP, BacMam 2.0) for overnight. After 24 hrs of the transfection, cells were fixed and processed for immnuostaining.

### Immunofluorescence and antibodies

Cells were fixed with 4 % paraformaldehyde in 1XPBS for 10 mins, permeabilized with 0.1 % Triton X-100 for 10 mins and blocked with 5 % BSA in PBS for 1 hr at room temperature. Primary antibodies were diluted in 5 % BSA in 1XPBS and slides were incubated overnight at 4°C. Slides were then rinsed thrice with 1XPBST(0.1% tween-20) and incubated with secondary antibodies for 1.5 hrs at room temperature. Slides were washed again and mounted with coverglass using Prolong Diamond Antifade Reagent with DAPI (Invitrogen; P36962). Images were obtained with a LSM 710 (Zeiss, X63, 1.4 NA) confocal microscope or Nikon Crest X-light 2 spinning disc confocal microscope (Nikon). Co-localization was analyzed with co-localization tool of ZEN software (Carl Zeiss). Primary antibodies used are FLAG (Sigma and Proteintech, 1:200), V5 (Life Technology, 1:200), PINK1 & TOM20 (Cell Signaling and Santa Cruz Biotechnology, 1:250), LC3B (Cell Signaling Technology, 1:250).

### Mitochondrial respiratory activity assay

Mitochondrial respiration in HeLa, HeLa^Parkin-YFP^ cells were measured using the Seahorse Extracellular Flux Analyzer XFp (Agilent Technologies, #102416-100) with XF Cell Mito Stress Test Kit (Agilent Technologies, #103015-100). Approximately, 1 x 10^4^ transfected cells were plated into desired drug-containing media in V3-PS 96 well plate the day before performing an assay. Next day, media were replaced with fresh assay media supplemented with 1 mM pyruvate, 2 mM glutamine and 15 mM glucose with a pH adjusted to 7.6. The standard mitochondria stress tests were performed consisting of basal measurements followed by measurements after sequential addition of 1 uM oligomycin, 0.5 uM FCCP and 0.5 uM Rotenone/antimycin A. At the end of the assay, protein concentration of each well were determined using BCA assay to normalize the obtained OCR values.

### Generation of *Drosophila* lines

Transgenic *Drosophila* lines were generated by BestGene with the standard PhiC31 integrase-mediated transgenesis system. All transgenes were inserted into the third chromosome attp2 site to avoid positional gene expression differences. Fly cultures and crosses were performed on standard fly food (Genesee Scientific) and raised at 25°C with a 12:12 hr light: dark cycle. MHC-GAL4 was used as driver for the expression in the muscles.

### Drosophila adult muscle preparation and immunohistochemistry

The sarcomere structure and mitochondrial morphology of the indirect flight muscle were analyzed as previously described(Baek et al., 2021). Adult flies were fixed with 4% paraformaldehyde in 1X PBS for 1 hour after a quick dissection,then immersed in OCT compound (Fisher Scientific) and flash frozen in liquid nitrogen or dry ice. Fixed tissues (∼15-16 microns) were sectioned by a cryomicrotome (Leica) and directly mounted on the slide. Additional post fixing was performed with 4% paraformaldehyde in 1XPBS for 10 mins at room temperature and then permeabilized with 0.2% Triton X-100 in 1XPBS and blocked with 5% BSA solution for 1 hr. Sections were incubated with phalloidin– Alexa Fluor 594 (Invitrogen) overnight at 4°C for muscle staining and mounted with Prolong Diamond Antifade Reagent with DAPI. Images were obtained with LSM 710 confocal microscope (Carl Zeiss) with 63X magnification.

### Drosophila ATP assay

Fly thoraces were dissected out, collected and homogenized in 20 µl of homogenization buffer (100 mM Tris, 4 mM EDTA and 6 M guanidine-HCL, pH 7.8). The homogenate was centrifuged at 16,000 x g for 10 min and the supernatant was collected and diluted to 1:200 with deiodinized water for ATP measurement using the Cell Titer-Glo luminescent cell viability assay kit (Promega, G7571) and 1:10 for determining the protein concentration. Data was obtained by normalizing the ATP concentration with protein amount.

### Image analysis and statistical analysis

Mitochondrial length was measured using MiNA tool set combined with ImageJ software as previously described(Valente et al., 2017). Statistical analysis was performed with Prism5(GraphPad) software. Statistical significance is expressed as *p* value that was determined with student’s *t*-test, one-way or two-way analysis of variance. One- or two-way ANOVA was followed by the indicated *post-hoc* tests.

## Acknowledgment

We thank Maddie Chalmers for her valuable feedback during manuscript preparation.

## Authors contribution

**Swati Maitra:** Performed experiments; analysed the data; prepared the manuscript. **Minwoo Baek:** Designed and performed the experiments; analysed the data. **Yun Jeong Choe:** Designed and performed experiments; **Nam Chul Kim:** Concieved and supervised and funded the project; reviewed and edited the manuscript.

## Funding

This research was supported by grants from NIA/NINDS **R56NS112296** (to Nam Chul Kim).

### Conflicts of Interest

The authors have no conflicts of interest to disclose.

**Correspondence to:** Nam Chul Kim

---

**Supplementary Figure S1.**
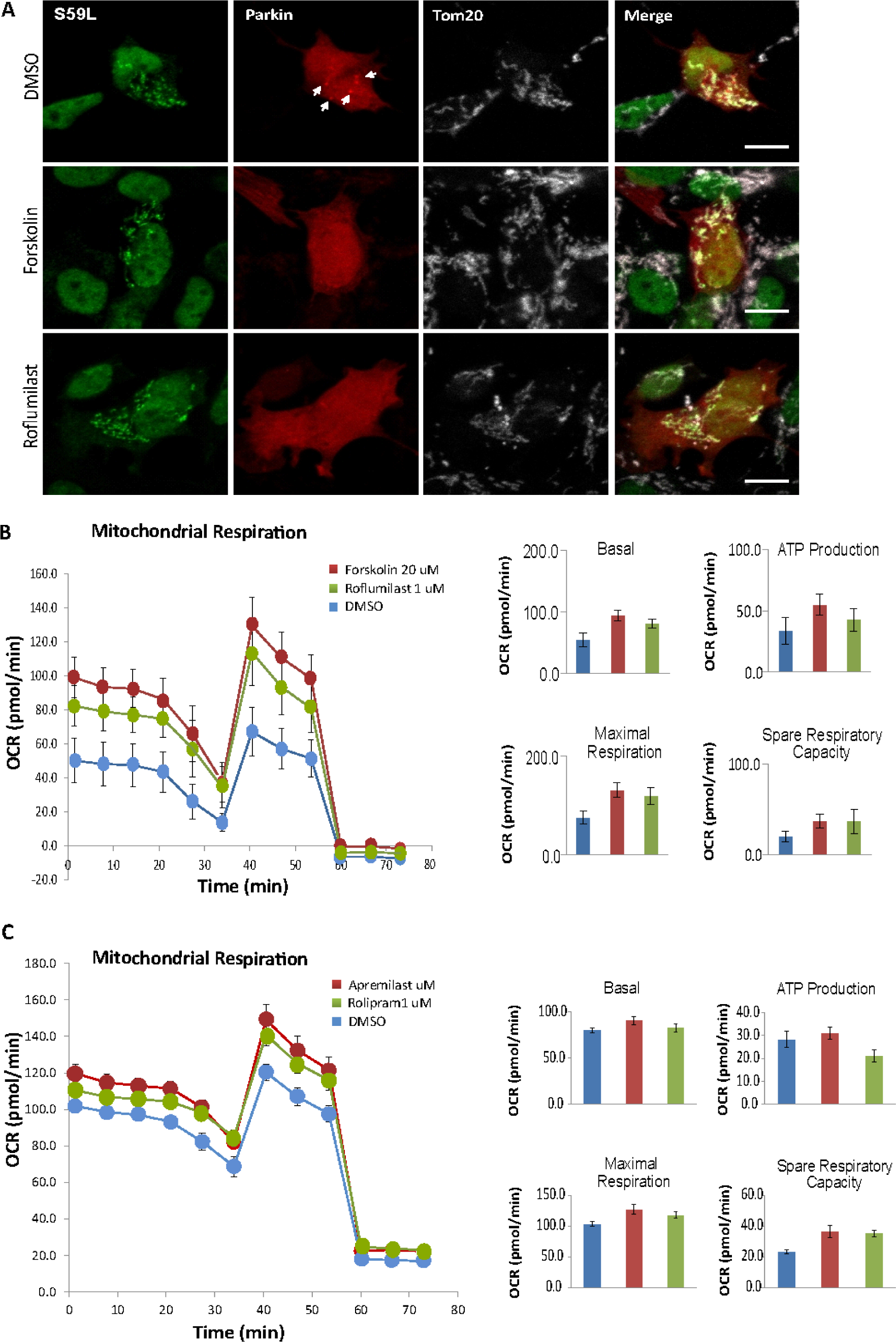
SH-SY5Y cells were co-transfected with FLAG-tagged CHCHD10^S59L^ and Parkin-mCherry in DMSO, Forskolin, and Roflumilast containing medium. **(A)** Representative images from CHCHD10^S59L^ expressing mitochondria of SHSY5Y cells cultured in DMSO, Forskolin and Roflumilast containing media. Cells were immunostained against anti-FLAG (green, CHCHD10^S59L^) and anti-TOM20 (white, mitochondria) antibodies. Arrows indicated accumulated parkin in the CHCHD10^S59L^ expressing mitochondria. **(B)** Mitochondrial respiration was measured by Seahorse XF Cell Mito Stress tests 24hrs after CHCHD10^S59L^ transfection in DMSO, Forskolin and Roflumilast treated SHSY5Y cells. **(C)** Mitochondrial respiration by Seahorse XF Cell Mito Stress tests 24hrs after CHCHD10^S59L^ transfection in DMSO, Apremilast and Rolipram treated SHSY5Y cells.

**Supplementary Figure S2.**
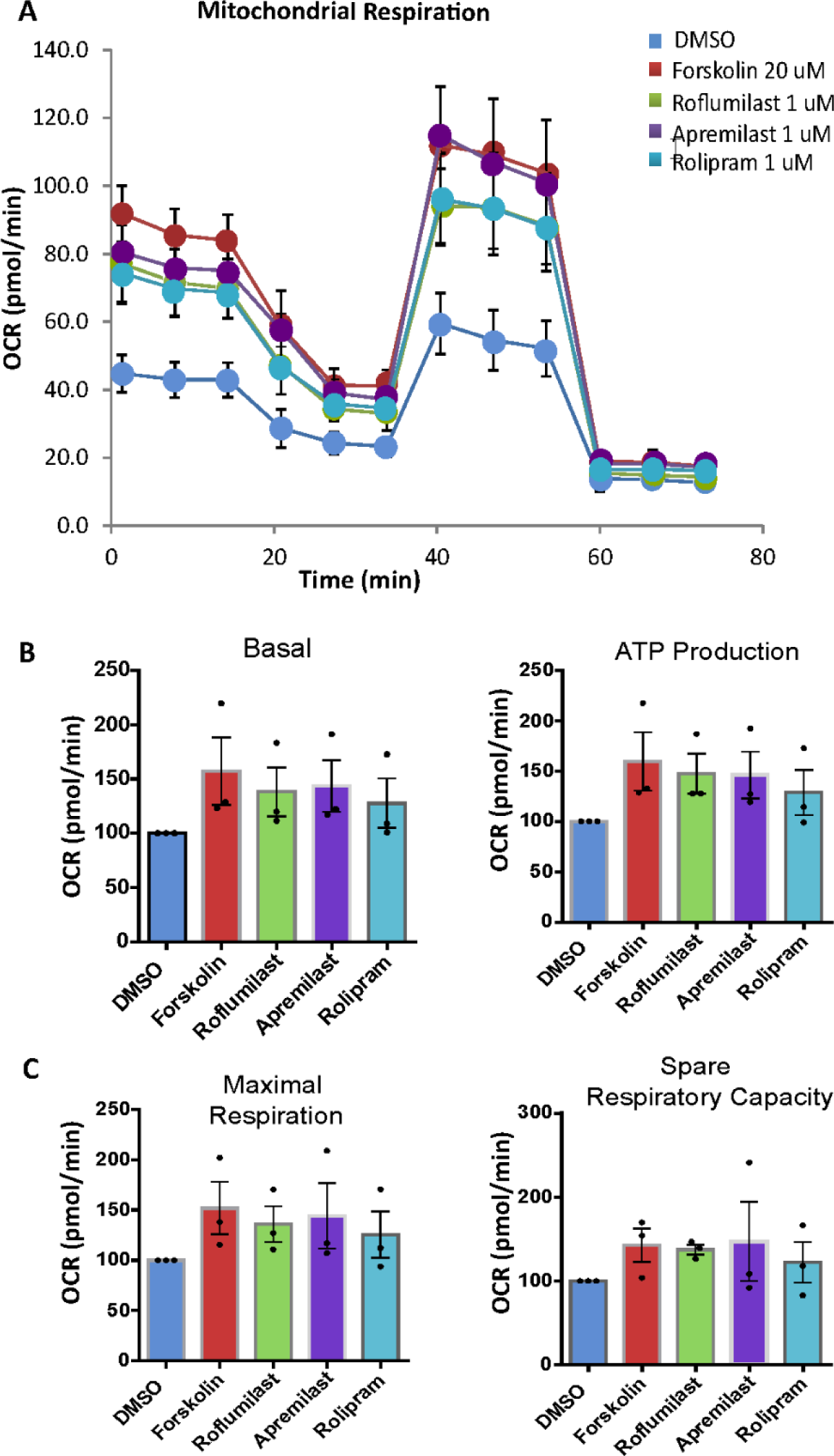
**(A)** Mitochondrial respiration was measurement by Seahorse XF Cell Mito Stress tests 24hrs after CHCHD10^S59L^ transfection in DMSO, Forskolin, Roflumilast, Apremilast and Rolipram treated HeLa cells. **(B) (C)** Bar graphs indicating basal respiration, ATP production, Maximal respiration, Spare respiratory capacity measured during seahorse assay in HeLa cells after CHCHD10^S59L^ transfection in DMSO, Forskolin, Roflumilast, Apremilast and Rolipram treatment. Data are shown as mean ±SD (one-way ANOVA and post hoc Dunnett test, two-sided, comparison with DMSO, \*\*\**p* < 0.001, \*\*\**p* < 0.01 and \**p* < 0.05).

